# Comparative genomic analyses highlight the contribution of pseudogenized protein-coding genes to human lincRNAs

**DOI:** 10.1101/163626

**Authors:** Wan-Hsin Liu, Zing Tsung-Yeh Tsai, Huai-Kuang Tsai

**Affiliations:** Institute of Information Science, Academia Sinica, Taipei, 115, Taiwan; Bioinformatics Program, Taiwan International Graduate Program, Academia Sinica, Taipei, 115, Taiwan; Institute of Bioinformatics and Systems Biology, National Chiao Tung University, Hsinchu, 300, Taiwan; Department of Plant Biology, Michigan State University, East Lansing, MI 48824, USA

**Keywords:** long intergenic noncoding RNAs, pseudogenization, transposable element, competing endogenous RNA, syntenic analysis

## Abstract

**Background:** The regulatory roles of long intergenic noncoding RNAs (lincRNAs) in humans have been revealed through the use of advanced sequencing technology. Recently, three possible scenarios of lincRNA origin have been proposed: *de novo* origination from intergenic regions, duplication from long noncoding RNA, and pseudogenization from protein. The first two scenarios are largely studied and supported, yet few studies focused on the evolution from pseudo genized protein-coding sequence to lincRNA. Due to the non-mutually exclusive nature that these three scenarios have, accompanied by the need of systematic investigation of lincRNA origination, we conduct a comparative genomics study to investigate the evolution of human lincRNAs.

**Results:** Combining with syntenic analysis and stringent Blastn *e*-value cutoff, we found that the majority of lincRNAs are aligned to the intergenic regions of other species. Interestingly, 193 human lincRNAs could have protein-coding orthologs in at least two of nine vertebrates. Transposable elements in these conserved regions in human genome are much less than expectation. Moreover, 19% of these lincRNAs have overlaps with or are close to pseudogenes in the human genome.

**Conclusions:** We suggest that a notable portion of lincRNAs could be derived from pseudogenized protein-coding genes. Furthermore, based on our computational analysis, we hypothesize that a subset of these lincRNAs could have potential to regulate their paralogs by functioning as competing endogenous RNAs. Our results provide evolutionary evidence of the relationship between human lincRNAs and protein-coding genes.

## Background

Long intergenic non-coding RNAs (lincRNAs) are a subclass of non-coding RNAs, which are longer than 200 nucleotides and locate between protein-coding genes. The advance of sequencing technology has recently revealed that lincRNAs present in various aspects of transcriptome and largely transcribed in many species from invertebrates to humans [1-3]. LincRNAs participate the regulation of many biological processes, such as the development of neuron components [3], gene expression [4, 5], and carcinogenesis [6]. Although the functions of lincRNAs are gradually explored, the originating mechanism of lincRNA has not attained to a conclusion. As the origination of lincRNAs would increase the regulatory complexity and impact several biological processes, it is intriguing to know where they evolve from and what the evolutionary mechanisms are.

Currently, three non-mutually exclusive possible mechanisms of lincRNAs origin have been proposed [1, 2, 7]. First, lincRNAs could be evolved from the duplication of other long non-coding RNAs (lncRNAs). Second, lincRNAs could be evolved from *de novo* origin, where the sequences could be previously noncoding or derived from transposable elements (TEs). Third, lincRNAs are considered to be originated from pseudogenization of protein-coding genes sequences. A well-studied lincRNA, *Xist*, which plays an important role in X chromosome inactivation [13, 14], is regarded as originated from protein-coding genes, *Lnx3*, by shown to contain the debris of a protein-coding genes sequences [15].

TE has been suggested as one of the major driving forces of lincRNA evolution. The capability of TE to transfer sequence to different regions across the genome allows new transcripts by providing valid regulatory sequences. Regulatory sequences such as promoter, transcription start site, enhancer, and splicing site can lead to transcribe a novel RNA [8] or process a precursor RNA into a stable transcript [9, 10]. A recent study has estimated about 10% of human lncRNA transcripts that were originated from long terminal repeats [8]. Moreover, many mature lncRNAs have been found entirely composed of endogenous retroviral sequences [8]. In addition to introducing regulatory sequence to cause lncRNAs, studies also found a substantial fraction of lncRNAs contains TE-derived sequences [11, 12], indicating a close association between TE and lncRNA.

However, due to the less conservation of lincRNAs sequences comparing to the mRNAs sequences, the studies of the lincRNAs from pseudogenization are more limited than the other two. In addition, the high TEs composition of exonic regions in lincRNAs (*i.e*. TEs inserted in the pseudogenes before and/or after the birth of the lincRNA) could also lead to underestimate the contribution of protein-coding gene pseudogenization to lincRNA origination [8]. Therefore, a detailed investigation of the lincRNAs originated by protein-coding gene pseudogenization is needed. Two key questions of what extents of the pseudogenized protein-coding genes contribute to lincRNA origination and whether there are some resulting regulations of these lincRNAs should be addressed.

The pseudogene has two different types, i.e. duplicated pseudogene and unitary pseudogene, that originate by different mechanisms and have distinct characteristic features [16]. A duplicated pseudogene was a copy of a gene which has been modified during and/or after duplication that resulted in loss of gene function. Alternatively, unitary pseudogene was a gene becoming disabled instead of a disable copy of a gene. Thus, the originating lincRNAs from protein-coding gene pseudogenization might follow such two different evolutionary trajectories, which has not been investigated thoroughly so far.

In this study, we focused on the human lincRNAs that might derive from protein-coding gene sequences. The potential origination for each human lincRNA was investigated by identifying its homologous sequences across nine vertebrate species. We analyzed sequence composition and genomic location of these putative orthologs to understand where human lincRNAs may originate from and what biogenesis components could contribute to the origination. According to our results, most of human lincRNAs have putative orthologs in at least two other vertebrates. Interestingly, although the majority locates in intergenic regions as expected, certain portions of these putative orthologs are partially or even fully annotated as protein-coding regions in at least two of the nine vertebrate species. We also found a subset of lincRNAs has conserved sequences in intronic regions in other species and the contribution of TEs to these alignments between lincRNAs and intronic sequences are marginal. To further explore the contribution of pseudogenized protein-coding gene to human lincRNAs, we determined which type of protein pseudogenization a lincRNA may originate from by investigate whether any human ortholog of its exonic ortholog in other species exists.

## Methods

### Genome annotation and sequence collection

First of all, cDNA sequences and genome coordinates of all the 7,340 human lincRNAs annotated in the Ensembl database (release 74) were downloaded [17]. These annotated lincRNAs were identified based on chromatin features and evidently low coding potential (*i.e*. for each lincRNA, no any known protein domain is found, and the predicted open-reading frame, if exists, is shorter than 35% of the total length). We further removed lincRNAs which overlap with human protein-coding genes to avoid potential bias in identifying putative orthologs. As a result, a total number of 6,618 lincRNAs were used in the following analysis.

Protein-coding sequences and genome annotations including non-coding gene annotations of the following nine vertebrate species were also downloaded from the Ensembl database: chimpanzee (*Pan troglodytes*; CHIMP2.1.4), orangutan (*Pongo abelii*; PPYG2), macaque (*Macaca mulatta*; MMUL_1), cow (*Bos taurus*; UMD3.1), dog (*Canis familiaris*; CanFam3.1), mouse (*Mus musculus*; GRCm38.p2), opossum (*Monodelphis domestica*; BROADO5), chicken (*Gallus gallus*; Galgal4), and zebrafish (*Danio rerio*; Zv9).

### Identification of putative orthologs of human lincRNAs in nine vertebrate species

We applied Blastn to identify matched sequences for cDNA sequences of each human lincRNA in the nine genomes. The parameters of Blastn (word size = 7, reward = 1, penalty = −1, and e-value < 10^−10^) were customized to increase the sensitivity for short alignment [18]. For each Blastn match, we then performed synteny analysis, which considers the order of conserved genes within the DNA regions between two species and has been shown to increase the reliability of identification of lincRNA homology region [19, 20]. We constrained at least one pair of conserved neighbor genes (*i.e*. one upstream and one downstream) to exist within ±750kb. We denote each of these regions as a candidate of putative ortholog (see a sketch map shown in **Fig. 1** and workflow in **Fig. S1**).

**Fig. 1.**
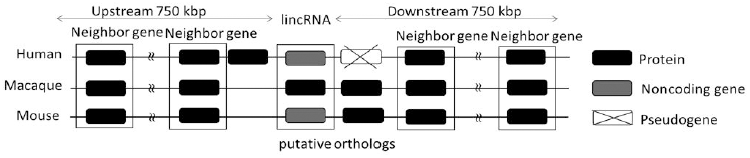
A sketch map illustrates the putative protein-coding orthologs of human lincRNAs in other species.

To further increase the confidence of our identification, any candidate of putative ortholog identified only in one of the nine species was removed. If there were multiple potential putative orthologs, we chose the one that was present most across the nine species. In the case of a continued tie, we selected the match with the lowest e-value. If a putative ortholog overlaps with at least one protein-coding gene, we annotated the putative ortholog as a protein-coding ortholog. If a putative ortholog did not overlap with a protein-coding gene but with at least one non-coding gene (*i.e*. either a lncRNA or a short non-coding RNA), we annotated the putative ortholog as a non-coding ortholog. The remaining putative orthologs were annotated as intergenic orthologs. To explore whether the human long non-coding sequences are associated with the exonic regions of protein-coding genes in other vertebrates, for each protein-coding ortholog, we calculated the percentages covered by exonic, intronic, and intergenic regions of protein-coding gene, respectively. The *exon/intron/intergenic coverage* was defined as the ratio between the length of exon/intron/intergenic-covered regions in the putative orthologs and the length of putative orthologs.

### Investigation of TEs in putative ortholog

To determine the coverage of TEs for each identified putative orthologs, we adopted RepeatMasker 4.0.3. [21], which was downloaded from Repeat Masker website (http://www.repeatmasker.org/) to identify TEs following the criteria used by Kapusta *et al*. [8]: only the sequences covered by more than 10 bps of RepeatMasker-annotated TEs were regarded as those derived from TE fragments. Furthermore, to study whether the annotated exon/intron/intergenic-covered regions were originated from TEs, we calculated *TE coverage* which was defined as the percentage of sequence identified as TEs by RepeatMasker.

### Examination of potential ceRNA role of lincRNA

According to the competing endogenous RNAs (ceRNA) theory [22], a RNA transcript that has microRNA binding site can sequester microRNAs from other RNA transcripts sharing the same microRNA binding site, thus regulating their expressions. Putative ceRNA-mRNA pairs annotated in the lnCeDB database [23] were used to examine whether lincRNAs could have potential to be ceRNAs for their putative paralog [23]. We assessed statistical significance of the observed number of ceRNA-mRNA pairs *N*_*obv*_ in the lincRNA-putative paralog pairs by using randomization via bootstrapping. Each time, we sampled *n* lincRNA-protein-coding genes from Ensembl database, where *n* is equal to the number of lincRNA-putative paralog pairs we found. The distribution of *N* was estimated given the null hypothesis that the number of ceRNA-mRNA pairs in the lincRNA-putative paralog pairs is the result of pure chance. For a one-tailed test with a rejection region in the upper tail, the bootstrap *p*-value *P(N_obv_)* for *N*_*obv*_, was estimated by the proportion of randomized samples that contain number of ceRNA-mRNA pairs > *N*_*obv*_. For *B* randomized datasets, we calculated the bootstrap *p*-value 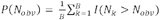 where *I(x)* is an indicator function yielding **1** if the ceRNA-mRNA pairs in *k-th* random dataset (*N*_*k*_) is more than in the original lincRNA-putative paralog pairs (*N*_*obv*_) and **0** otherwise. Here we conducted a bootstrapping analysis with *B* = 10000.

### Bootstrapping analysis of homologous sequences

A bootstrap analysis was performed to prove that the homologous sequences identified in our study are not simply artifact. By focusing on the 193 lincRNAs having protein-coding orthologs, we first shuffled each of the sequences 5,000 times with control of di-nucleotide content. We then performed Blastn (with the same setting: word size = 7, reward = 1, penalty = −1) to align shuffled sequences to syntenic regions (with the same definition: −750 kb to +750 kb sequence with conserved order of orthologous neighbor genes) if any in the nine vertebrate species. For each shuffled sequence, we selected the hit with the lowest Blastn *e*-value as the alignment. According to our shuffling analysis, none of the shuffled sequences can have a hit with blast *e*-value < 10^−10^. The Blastn *e*-value distributions of shuffled sequences and original lincRNAs are shown in Fig. S2. The results show that our method of syntenic analysis and Blastn *e*-value is sufficient to discriminate real homologous relationships from noise.

## Results and Discussion

### De novo origination and protein pseudogenization contribute to lincRNA evolution mostly

We performed syntenic analysis with nine vertebrate genomes to identify putative orthologs of human lincRNAs (see Method). A putative ortholog was defined as a region that has significant sequence similarity and share the same synteny with human lincRNAs (**Fig. 1**). Based on the genome annotations, putative orthologs were further classified into three groups: putative intergenic orthologs, putative protein-coding orthologs, and putative non-coding orthologs (*Table S1*, see Methods). According to the proportion of each human lincRNA group in the nine species, the majority of their putative orthologs belonged to intergenic orthologs, followed by protein-coding orthologs, and only few putative orthologs were classified as non-coding orthologs (**Fig. 2**). This result reflects the relative contributions of the three lincRNA origination scenarios: *de novo* origination, protein pseudogenization, and duplication from lncRNA.

**Fig. 2.**
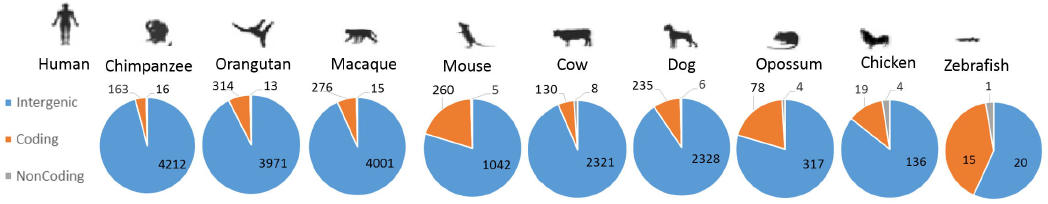
The ratios of genomic loci of putative orthologs of human lincRNAs in nine species. Most lincRNAs have putative orthologs being annotated as intergenic regions (4,212 in chimpanzee, 3,971 in orangutan, 4,001 in macaque, 2,321 in cow, 2,328 in dog, 1,042 in mouse, 317 in opossum, 136 in chicken, and 20 in zebrafish). Nevertheless, for a notable number of lincRNAs, the corresponding putative orthologs are overlapping with protein-coding genes (the numbers of coding orthologs as defined in Methods are: 163 in chimpanzee, 314 in orangutan, 276 in macaque, 130 in cow, 235 in dog, 260 in mouse, 78 in opossum, 19 in chicken, and 15 in zebrafish). The corresponding putative orthologs overlapping with non-coding genes are less comparing to intergenic regions and protein-coding gene (16 in chimpanzee, 13 in orangutan, 15 in macaque, 8 in cow, 6 in dog, 5 in mouse, 4 in opossum, 4 in chicken, and 1 in zebrafish).

The predominant number of putative intergenic orthologs could be mainly contributed by *de novo* origination, in particular the insertion of TEs. TEs have been found to be abundant in intergenic regions due to their moving and amplifying ability [8, 24-26]. Together with the high TEs composition of lincRNA [8, 27, 28], our observation supports that TE is one of the major factors contributing to lincRNA origination. Another possible explanation of intergenic orthologs might be incomplete annotation of non-coding RNAs. Moreover, current annotation of lincRNA could bias to primates [29], thus lead to underestimate the contribution of lncRNA duplication in lincRNA origination.

As the second large group in the putative orthologs, although the ratio varies significantly across species (e.g. 4% in chimpanzee and 40% in zebrafish), the numbers of putative protein-coding orthologs are around 160 to 310 in the closer species. Among the total 6,618 human lincRNAs, 297 of them (4.5%) have protein-coding orthologs in at least two vertebrates. Our results indicate certain contribution of protein-coding gene pseudogenization to human lincRNAs as reported in the previous study which a lincRNA (*e.g. Xist*) can retain both the syntenic context and the debris of the exon of protein (*e.g. Lnx3*) [15]. Although the number is less than the number of putative intergenic orthologs, the amount of putative protein-coding orthologs is more than previously documented. The reason why the number of putative protein-coding orthologs is more than previously documented could be in previous studies, to found the lncRNA orthologs and to avoid the high coding potential bias of the non-coding RNA sequences, the RNAs that have high coding potential such as mRNAs and pseudogenes are often kicked out in the lncRNA datasets in early parsing process.

The evidence of protein pseudogenization is stronger when there are annotated orthologous relationships among the genes where a putative ortholog resides in different species (referred as aligned proteins). Hence, we adopted orthologous relationships in Ensembl [30] and show that 193 of 297 (65%) of the aligned proteins are annotated as orthologous pairs. For example, human lincRNA AC004471.10 possesses aligned proteins TSSK2 in cow, ENSCAFG00000023784 in dog, ENSMMUG00000031114 in macaque, and Tssk2 in mouse, which are orthologous with each other. To sum up, the significantly greater numbers of intergenic and protein-coding orthologs than non-coding orthologs reveal the important roles of *de novo* origination and protein pseudogenization in lincRNA evolution.

Lastly, the number of non-coding orthologs here was much less than the others. One explanation is, as mentioned previously, the incomplete and biased annotation of non-coding RNAs on other species. Another reason is the relative small number of non-coding genes in the reference genome and the poor conservation of lincRNAs. The number of non-coding genes in each reference species is around 10%-30% of protein-coding genes. Because protein-coding orthologs only account for 30% of putative orthologs or less, the numbers of non-coding orthologs are less than 3%. In addition, we could not identify any non-coding orthologs of lincRNA in mouse or zebrafish genome. The result agreed with recent study in zebrafish that reported merely a minority of lincRNAs showed significant sequence similarity to other lncRNAs [18].

### TEs have only minor contribution in the sequence similarity between a lincRNA and its protein-coding ortholog

Since the conserved introns have been proposed to be a potential source of lncRNA [31], the corresponding orthologs of the lncRNAs locating within an open reading frame (ORF) may not be originated from protein pseudogenization. The investigation in the gene structure (*i.e*. the coverages and distributions of exons and introns) of protein-coding orthologs is needed to distinguish if protein pseudogenization was involved in lincRNA evolution. To evaluate how many protein-coding orthologs could be considered as evidences of protein pseudogenization, *exon coverage*, *intron coverage,* and *intergenic coverage* were examined for each protein-coding ortholog (see Methods).

The distributions of exon coverage, intron coverage, and intergenic coverage for each protein-coding ortholog are shown in **Fig. 3**. Considering all of the putative orthologs from the nine vertebrates, the results showed 692 putative protein-coding orthologs (32%) in which *exon coverage* was greater than both *intron coverage* and *intergenic coverage*. Moreover, 263 (12%) fully located within exonic regions (*i.e. exon coverage =* 100%), which were defined as exonic orthologs in this study. On the contrary, 1124 (52%) putative protein-coding orthologs were intronic orthologs, which completely located within intronic regions (*i.e. intron coverage =* 100%). Studies have suggested that some lncRNAs could be post-processed into small nucleolar RNAs (snoRNAs) [32], which are involved in ribosome synthesis or translation, and are usually intronic sequences [33]. The hypothesis is that lncRNAs could be post-processed into snoRNAs and involved in ribosome synthesis and translation mechanism. Therefore, one of the possible explanations is that these conserved intronic orthologs might be unannotated lncRNAs.

**Fig. 3.**
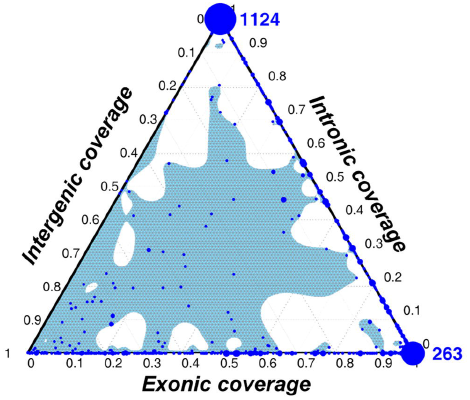
Ternary plot of *intronic coverage, intergenic coverage*, and *exonic coverage*. 1124 (52%) protein-coding orthologs that are completely intronic. Alternatively, 263 (12%) are completely exonic. The size of each dot correlates with the number of lincRNAs having this combination of *exonic coverages, intronic coverages,* and *intergenic coverages*. The blue shadow illustrates the estimated distribution.

Besides, TEs could be an alternative explanation for the high intronic coverage because TEs are known to locate in introns more than in exons [8, 34], and a high TE composition are reported in lincRNAs [8, 27, 28]. Thus, we identified TE for each putative protein-coding ortholog using RepeatMasker and calculated *TE coverag*e for each putative protein-coding ortholog (see Methods). The results showed that TEs covered 47% region when considering all introns of protein-coding orthologs jointly. Unexpectedly, low *TE coverages* (**Fig. 4(a)**, average = 0.32) were observed even the intronic orthologs. Similarly, *TE coverages* of exonic orthologs and intergenic orthologs were also low (**Fig. 4 (b) and 4(c)**, average = 0.07 and 0, respectively). Taking together, insertion of TEs may only contribute to a minor part of the sequence similarity between lincRNA and protein-coding orthologs.

**Fig. 4.**
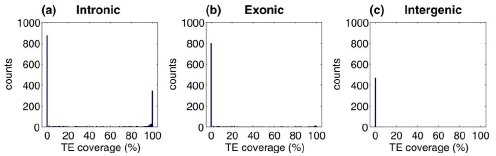
The distributions of *TE coverages* in the putative orthologs of lincRNA. (a) intronic orthologs, (b) exonic orthologs, and (c) intergenic orthologs.

### LincRNAs derived from duplicated pseudogenes could impact the regulation of their putative paralogs

Exonic orthologs of lincRNAs identified in our study showed that a certain portion of human lincRNAs could be derived from protein pseudogenization (**Fig. 3**). According to extremely low *TE coverages* in the protein-coding orthologs (Fig. 4), the results implied the minor contribution of TEs in origination of lincRNA. In particular, the *TE coverage*s were zero in the exonic orthologs of 108 lincRNAs (*i.e*. without TE insertion). Consequently, we ask what mechanism causes such protein pseudogenization that involved in lincRNA origination.

GENCODE project [16] has categorized pseudogenes into three main groups based on genomic features and evolutionary mechanisms: processed pseudogenes, duplicated (also referred to as unprocessed) pseudogenes, and unitary pseudogenes. Processed pseudogenes were originated from retrotransposition which was a fraction of mRNA back into the genome since they contain only exonic sequence and do not retain the upstream regulatory regions. In contrast, duplicated pseudogenes have intron-exon like genomic structures and may still maintain the upstream regulatory sequences of their parents, as a result, duplicated pseudogenes might derive from duplication of functional genes. Last, since unitary pseudogene lost their coding parental gene in human and can only find the coding ortholog in a reference species, unitary pseudogene might be originated from accumulated fixed disabling mutations in a coding gene [16].

Among the 108 lincRNAs having exonic orthologs, 66 of them do not have any putative paralog nor TE coverage. Moreover, sixteen (24.2%) of them overlapped with pseudogenes and eight (12.1%) of them are just next to pseudogene, suggesting the potential origination from unitary pseudogene. One example is lincRNA CTD-2555O16.1 which overlapped with pseudogene TEX21P, as shown in **Fig. 5**. On the other hand, 42 lincRNAs have putative paralogs, that is, their aligned proteins possess at least one human ortholog (e-value less than 10^−10^). In addition, among these 42 lincRNAs, 13 (31%) overlapped with known pseudogene. Take lincRNA RP5-998N21.4 for example (**Fig. 6**), its transcript overlapped with the transcript of pseudogene FCGR1C. Moreover, four lincRNAs such as CROCCP2 and ADAM20P1 are annotated as pseudogenes of human orthologs of their aligned proteins from the Ensembl records.

**Fig. 5.**
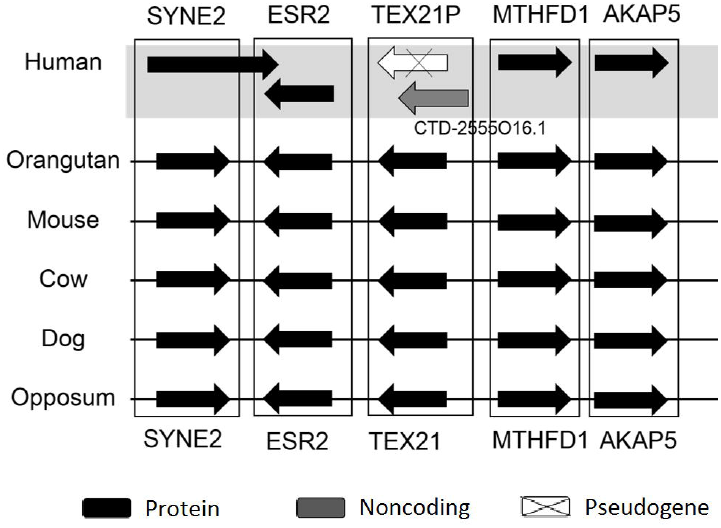
The syntenic regions across six species of lincRNA CTD-2555O16.1 which overlapped with unitary pseudogenes TEX21P. The human pseudogene TEX21P possesses homologous protein Tex21 in orangutan, mouse, cow, dog, and opossum. Protein-coding genes are indicated in black, RNA-genes in gray, and pseudogenes in white.

**Fig. 6.**
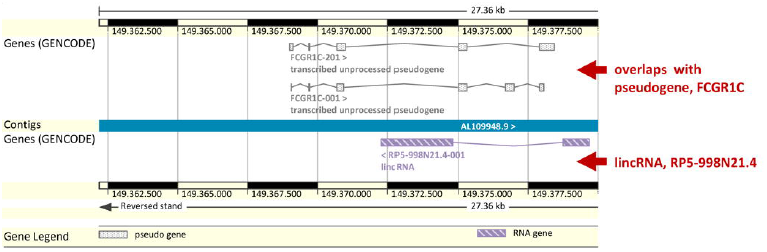
An example of lincRNAs overlapped with known pseudogenes. LincRNA RP5-998N21.4 overlapped with the transcripts of pseudogenes FCGR1C (figure modified from Ensembl genome browser Ver.74). As this particular lincRNA overlaps in antisense with the transcribed pseudogene, the regulatory sequences involved in the lincRNA expression could be different than the ones of the processed pseudogene.

A hypothesized explanation of such relationships is that lincRNAs regulate their putative paralog by functioning as a competing endogenous RNA (ceRNA). Based on the high sequence identity and close genomic positions between lincRNA and its putative paralog, it is possible that lincRNA regulates its putative paralog as a ceRNA. According to the annotation in the lnCeDB database [23], 28 lincRNA-putative paralog pairs were annotated as ceRNA-mRNA pairs. We further performed a bootstrapping analysis (see Methods) to test whether ceRNA-mRNA pairs annotated in lnCeDB database were enriches in these lincRNA-putative paralog pairs identified in the present study. The significant enrichment (bootstrap *p* < 6.8 × 10^−3^) supports the proposed hypothesis that lincRNA could possibly regulate its putative paralog.

Through Gene Ontology enrichment analysis, most of these putative paralogs were found to have binding functions (metal ion binding, *p* = 5.05 × 10^−9^; cation binding, *p* = 9.64 × 10^−9^; DNA binding, *p* = 1.12 × 10^−8^; nucleic acid binding, *p* = 1.88 × 10^−7^; heterocyclic compound binding, *p* = 1.80 × 10^−4^; organic cyclic compound binding, *p* = 2.65 ×10^−4^; ion binding, *p* =6.31 × 10^−4^). In addition, these proteins significantly associate with neuron development and eye disorder [35-37] according to the Online Mendelian Inheritance in Man (OMIM) database [38]. With conserved sequences, lincRNA could influence expression of putative paralogs by post-transcriptional regulation as endogenous siRNA or buffering effect as their decoys. Moreover, we cannot rule out the possibility that some genes are functional as well in RNA level thus their paralogous lincRNA could also contribute to these particular functions.

## Conclusions

Protein pseudogenization is one of the scenarios of human lincRNA originations. According to the comparative genomics analyses among human and nine vertebrate species in this study, 193 of the 6,614 human lincRNAs have protein-coding orthologs, which are conserved sequences in protein-coding genes of other species. Our study reveals the role of protein pseudogenization in human lincRNA origination. We anticipate that these results will bring insights to the evolutionary originations and genetic functionalities of human lincRNAs.

## Declarations

### List of abbreviations

lincRNA: long intergenic noncoding RNA
TE: transposable element
lncRNA: long non-coding RNA
ORF: open reading frame
ceRNA: competing endogenous RNA

## Ethics

Not applicable

## Consent to publish

Not applicable

## Authors’ contributions

WHL, ZTYT, and HKT designed the analyses. WHL collected the data and performed the analyses. WHL, ZTYT, and HKT wrote the paper. HKT was the principal investigator. All authors read and approved the final manuscript.

## Availability of data and materials

The datasets supporting the conclusions of this article are included within the article and its additional files.

## Acknowledgements

The authors are thankful to Jen-Hao Cheng, Wen-Yi Chu, and Jia-Hsin Huang for their suggestions on this manuscript.

## Additional files

**Additional file 1 - Table S1**. The putative orthologs of human lincRNAs in the nine analyzed species.

**Additional file 2 - Fig. S1**. The workflow for identifying the putative orthologs of lincRNAs in the nine species.

**Additional file 3 - Fig. S2**. The Blastn *e*-value distributions of shuffled sequences and 193 lincRNAs (see Methods).

